# LION/web: a web-based ontology enrichment tool for lipidomic data analysis

**DOI:** 10.1101/398040

**Authors:** Martijn R. Molenaar, Aike Jeucken, Tsjerk A. Wassenaar, Chris H. A. van de Lest, Jos F. Brouwers, J. Bernd Helms

## Abstract

A major challenge for lipidomic analyses is the handling of the large amounts of data and the translation of results to interpret the involvement of lipids in biological systems. We built a new lipid ontology (LION) that associates over 50,000 lipid species to biophysical, chemical and cell biological features. By making use of enrichment algorithms, we used LION to develop a web-based interface (LION/web, www.lipidontology.com) that allows identification of lipid-associated terms in lipidomes. LION/web was validated by analyzing a lipidomic dataset derived from well-characterized sub-cellular fractions of RAW 264.7 macrophages. Comparison of isolated plasma membranes with the microsomal fraction showed a significant enrichment of relevant LION-terms including ‘plasma membrane’, ‘headgroup with negative charge, ‘glycerophosphoserines’, ‘above average bilayer thickness’, and ‘below average lateral diffusion’. A second validation was performed by analyzing the membrane fluidity of CHO cells incubated with arachidonic acid. An increase in membrane fluidity was observed both experimentally by using pyrene decanoic acid and by using LION/web, showing significant enrichment of terms associated with high membrane fluidity ('above average’, 'very high’ and 'high lateral diffusion’, and 'below average transition temperature’). The results demonstrate the functionality of LION/web, which is freely accessible in a platform-independent way.

## MAIN TEXT

The comprehensive study of lipids, also termed lipidomics, is gaining momentum. Instrumentation is becoming increasingly more sensitive, precise and fast, and the use of lipidomics to address key questions in membrane biology has become widespread. As a result, datasets are rapidly increasing both in terms of size and complexity. Due to a lack of methods to perform global and in-depth data mining, lipidomic research tends to focus on individual lipid classes or lipid species. A common approach in other ‘omics’ disciplines to reduce complexity is the use of ontologies *e.g.*, Gene Ontology (Ashburner et al., 2000), Chemical Entities of Biological Interest ontology (Degtyarenko et al., 2008), combined with statistical tools to determine terms of interest.

Although lipid structure is closely related to lipid function, it is currently impossible to associate properties of individual lipids with complex lipid mixtures of cellular lipidomes. Examples of biophysical properties that play an important role in membrane biology are numerous and include membrane thickness as driving force in the sub-cellular localization of proteins (Sharpe et al., 2010), membrane fluidity regulating bacterial survival (Inda et al., 2014), membrane heterogeneity in cellular signaling (Sezgin et al., 2017), intrinsic curvature of lipids as key player in lipid droplet biogenesis (Ben M’barek et al., 2017; Thiam et al., 2013) or COPI coat disassembly (Bigay et al., 2003), and net charge of membranes as a determinant in lipid-protein interactions (Enkavi et al., 2017).

Here, we aim to provide a lipid ontology database and complementary enrichment analysis tool that (i) contains chemical and biophysical information of lipid species, (ii) is platform independent and compatible with routine mass spectrometry-based lipid analysis, (iii) can be used by researchers without computer programming skills, and (iv) is freely available to the scientific community.

We constructed an ontology database called LION that links over 50,000 lipid species with four major branches: ‘lipid classification’ (Fahy et al., 2009), ‘chemical and physical properties’ (fatty acid length and unsaturation, headgroup charge, intrinsic curvature, membrane fluidity, bilayer thickness), ‘function’, and ‘subcellular localization’ (as described in literature). The resulting database contains more than 250,000 connections (‘edges’), providing a detailed system for in-depth annotation of lipids. An example of all LION-terms associated with a single phosphatidylserine (PS) lipid species, PS(34:2), is depicted in **Figure S1**.

An important feature of LION is the association of lipid species with biophysical properties. We made use of experimental data (Marsh, 2010) and data obtained by coarse-grain molecular dynamics simulation (CG-MD) (Wassenaar et al., 2015), each providing distinct biophysical properties. These data were used to estimate the biophysical properties of all related lipids in the LION-database by multiple regression analysis.

The regression models were validated in two ways. First, we performed leave-one-out cross-validations (LOOCV) of all three models (**Fig. S2 A-C**), showing satisfactory agreement between determined and predicted values. Second, we compared two properties closely associated with membrane fluidity: ‘transition temperature’ (from experimental datasets) and ‘lateral diffusion’ (from the CG-MD datasets) (**Fig. S2 D**). As expected, lipids with low transition temperatures were predicted to have high lateral diffusion values at a defined simulation temperature and vice versa.

Subsequently, all numerical datapoints for each biophysical property were categorized into five pre-defined groups (‘very low’, ‘low’, ‘average’, ‘high’, ‘very high’). The limits of each group were determined based on the presence of lipid species reported in four lipidomics publications (Andreyev et al., 2010; Haraszti et al., 2016; Köberlin et al., 2015; Lin et al., 2017). These values were subsequently used to categorize all applicable lipid species present in LION (**Fig. S2 E**).

Next, we used LION as a basis to build an ontology enrichment tool that facilitates reduction of lipidome complexities in an unbiased manner. We made use of an adapted version of ‘topGO’, an R-package designed for enrichment analysis of GO-terms (Alexa and Rahnenfuhrer, 2017). Subsequently, we designed a web-tool with R-package Shiny (‘LION/web’, www.lipidontology.com) that offers an intuitive user-interface and supports two major workflows (**Fig. 1 and Note S1**): enrichment analysis of a subset of lipids of interest (‘by target list’) and enrichment analysis performed on a complete and ranked list of lipids (‘by ranking’, referred to as ‘SAFE’ in the context of genes (Barry et al., 2005)).

**Figure 1.**
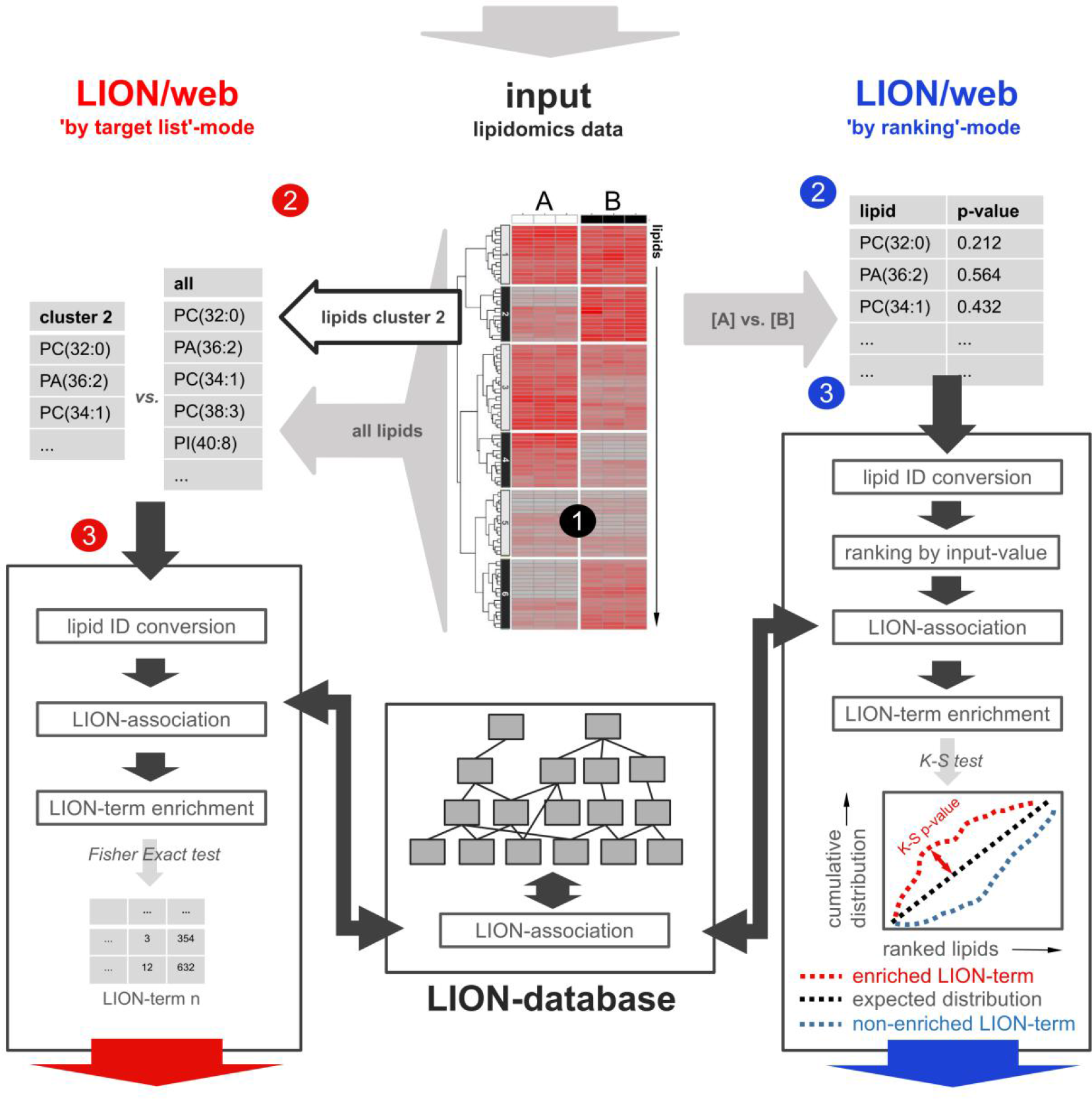
Enrichment analysis approaches supported by LION/web. A lipidomics dataset containing lipid identifiers and abundances derived from two or more conditions 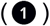 can be processed in two ways by LION/web. In the ‘by target list’-mode (left, 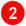), a subset of lipids (*e.g.*, derived from thresholding or clustering) is compared to the total set of lipids. After standardization of lipid nomenclature 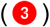, applicable LION-terms are associated and assessed for enrichment in the subset by Fisher’s exact statistics. Alternatively, in the ‘by ranking’-mode, input lipids are ranked by the provided values (‘local’ statistics). By default, *P* values from one-tailed t-tests are used 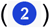. After ranking, lipid nomenclature is standardized 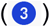. Applicable LION-terms are subsequently associated to the dataset and distributions are compared to a uniform distribution by ‘global’ statistics (here, Kolmogorov–Smirnov tests). Calculated *P* values of LION-terms from both approaches are corrected for multiple testing (Benjamini-Hochberg).

To test the functionality of LION/web, we made use of a previously published and well characterized dataset containing lipidomics data from several sub-cellular fractions of RAW 264.7 macrophages (Andreyev et al., 2010). First, we re-normalized the dataset by expressing all lipid species as fraction of the total amount of lipid per sample. LION/web was able to reformat (for information about input conventions, see **Note S2**) and match the vast majority (>97%) of the submitted lipids. Subsequently, we compared the isolated plasma membrane (PM) with the endoplasmic reticulum (ER) fraction from non-stimulated macrophages and assessed all LION-terms for enrichment (**Fig. S3**). In good agreement with current descriptions of the selected organelles (Holthuis and Menon, 2014; van Meer et al., 2008), significant enriched LION-terms included terms associated with chemical descriptions (*e.g.*, ‘glycerophosphoserines’, ‘headgroup with negative charge’, ‘phosphosphingolipids’), biological features (‘plasma membrane’) and biophysical properties (*e.g.*, ‘above average bilayer thickness’, ‘below average lateral diffusion’, ‘very low lateral diffusion’, ‘very high bilayer thickness’, ‘neutral intrinsic curvature’). LION/web also reported the significant enrichment of ‘very high transition temperature’, which is in line with the (very) low lateral diffusion terms (see also **Fig. S2 D**). Also the term ‘very low transition temperature’ was reported to be significantly enriched. Inspection of the lipid species responsible for the LION-term ‘very low transition temperature’ revealed the presence of lipids that all contain polyunsaturated fatty acids (PUFAs) with at least four unsaturations. This may be a macrophage-specific phenomenon, related to their involvement in inflammation (Calder, 2015).

To further validate LION/web, we used two different experimental approaches. First, we investigated the enrichment of LION-terms associated with chemical features that can be easily incorporated into lipids (*e.g.*, fatty acids as building blocks). To this end, CHO-k1 cells were incubated overnight in the presence of palmitic acid (PA), linoleic acid (LA) or arachidonic acid (AA) complexed to bovine serum albumin (BSA). Subsequently, lipids were analysed by LC-MS/MS (**Data S1**and **Fig. S4**)and submitted to LION/web (‘by ranking’-mode). LION/web offers the option to limit analysis to specific terms of interest. After pre-selection of LION-terms that indicate the presence of fatty acids as lipid building blocks, LION/web reported the significant enrichment of the respective fatty acid in the three different conditions (**Fig. 2 A** and **Data S2**).

**Figure 2.**
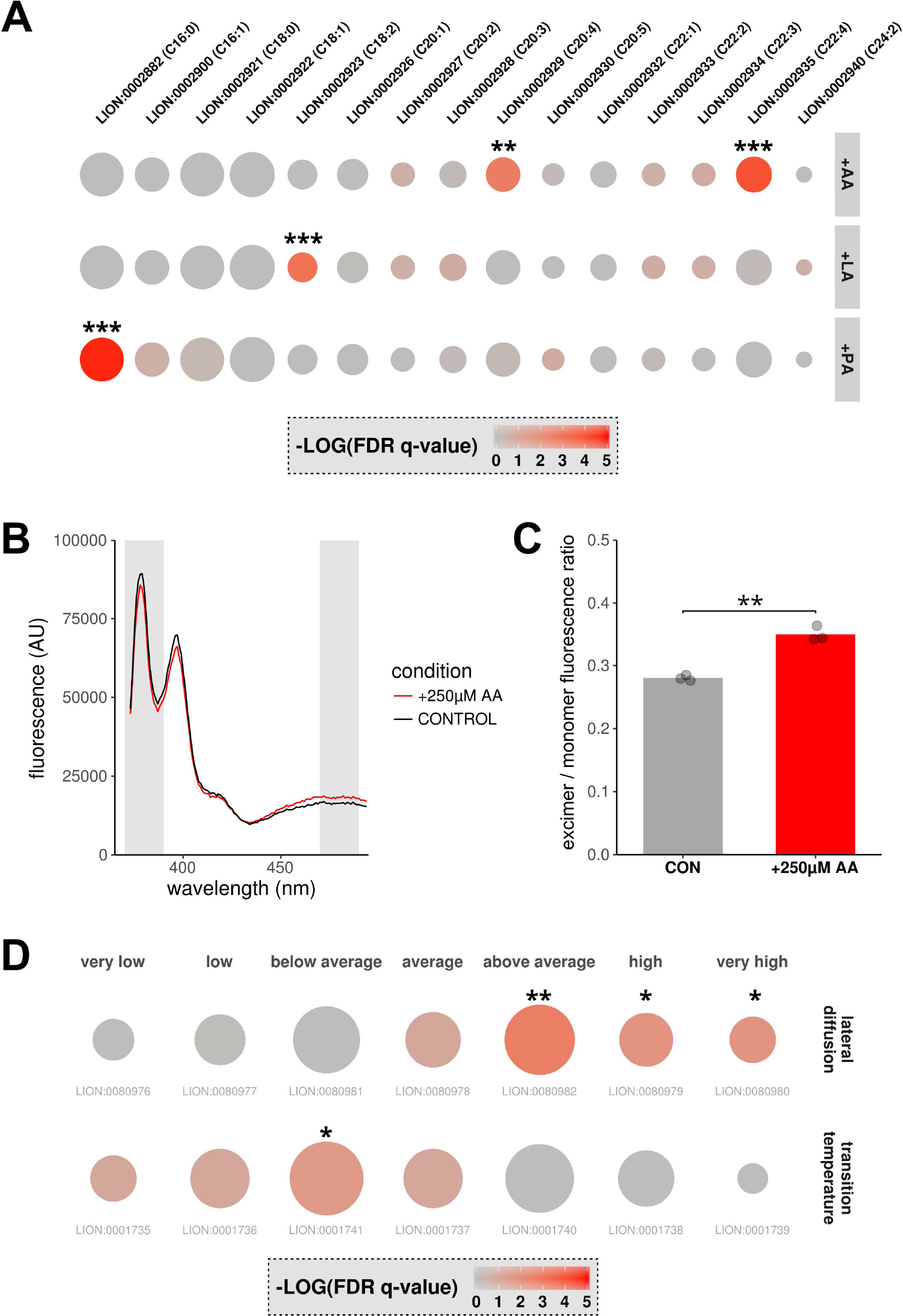
LION-term enrichment and membrane fluidity of CHO-k1 cells. CHO-k1 cells were incubated overnight with PA, LA or AA (100 μM, complexed to BSA) (**A**)or with AA (250 μM, complexed to BSA) (**B-D**). All incubations were performed in triplicate. For control incubations, cells were incubated with fatty-acid free BSA. (**A,D**)After extraction and lipidomics profiling by LC-MS/MS, enrichment analyses of the conditions of interest versus control incubations were performed by LION/web of (**A**)LION-terms indicating the presence of selected fatty acids or (**D**)LION-terms indicating the degree of membrane fluidity. Dot sizes in the dot plots are scaled to the number of associated lipids; colors are scaled to the level of enrichment. (**B,C**)After incubation, fluorescence emission spectra between 370 and 500 nm of lysates containing pyrenedecanoic acid (PDA) were measured in triplicate (**B**). Fluorescence spectra examples of either control (black) or AA-stimulated lysates (red). Gray shades indicate monomer and excimer fluorescence filters. (**C**)Mean ratios (bar) and individual datapoints (dots) of excimer over monomer fluorescence (representative data of three independent experiments). Statistical significance was determined by Student’s two-tailed t-test. (**A,C,D**)* *P* < 0.05, ** *P* < 0.01, *** *P* < 0.001.

Second, to investigate the enrichment of biophysical LION-terms, we incubated CHO-k1 cells with arachidonic acid (AA). This procedure is known to increase membrane fluidity (Yang et al., 2011). After incubation, the membrane fluidity properties of the samples were analyzed both experimentally and by LION/web. Membrane fluidity was experimentally assessed using pyrene decanoic acid (PDA). This fluorescent probe can exist as monomer or excimer, resulting in a shift of its emission spectrum. The ratio of excimer over monomer fluorescence is proportional to the degree of membrane fluidity (Eisinger and Scarlata, 1987). To this end, fluorescence spectra of lysates from cells incubated with or without AA were measured (**Fig. 2 B**). As expected, the ratio of excimer/monomer forms of PDA revealed a significant increase in membrane fluidity of lysates in the presence of AA (**Fig. 2 C**). For parallel LION/web analysis of membrane fluidity properties, lipids were extracted from the same samples and analysed by LC-MS/MS (**Data S3** and **Fig. S5**). LION contains two sets of terms associated with membrane fluidity: ‘transition temperature’ and ‘lateral diffusion’.

Accordingly, LION/web was set to limit enrichment analyses to these sets, after which the lipidomic data were analyzed (‘by ranking’ mode). In line with the experimentally measured increase in membrane fluidity, terms associated with high membrane fluidity ('above average’, 'very high’ and 'high lateral diffusion’, and 'below average transition temperature’) were significantly enriched in cells that had been treated with AA (**Fig. 2 D** and **Data S4**).

Taken together, we have presented a new ontology called LION that enables flexible annotation of lipid species and that covers most commonly found lipid classes and fatty acid distributions. Furthermore, it combines the well-established lipid class hierarchy from LIPIDMAPS with biophysical data that were not previously available. To explore lipid datasets in an unbiased manner, we built an online web-tool that does not require knowledge of programming languages. We believe that this lipid database and associated web-tool bridges the gap between lipidomics and cell biology by revealing patterns that are of biological interest.

## ACKNOWLEDGEMENTS

We thank Xin He, PhD, for providing and supporting the topONTO R-package. We thank Jeroen W.A. Jansen for the excellent technical assistance with the lipidomics experiments.

## AUTHOR CONTRIBUTIONS

M.R.M. and J.B.H. conceived the project. M.R.M. developed LION, LION/web and performed the experiments. A.J. tested and suggested improvements for LION/web. C.H.A.v.d.L. and T.A.W. contributed to the regression models and statistical concepts. C.H.A.v.d.L. and J.F.B. contributed to the lipidomics data processing and analysis. M.R.M. and J.B.H. wrote the manuscript.

## COMPETING FINANCIAL INTERESTS

The authors declare no competing financial interests.

## Methods

### Creation of lipid ontology (LION)

We built an ontology database that connects over 50,000 lipid species to the following four major branches: ‘lipid classification’, ‘function’, ‘cellular localization’ and ‘physical or chemical properties’. For readability, a term is included at the top of each branch to indicates the nature of a LION-branch. These ‘category’ terms are distinguished from other LION-terms with an ID containing the prefix ‘CAT’.

The classification system is based on the LIPIDMAPS classification (Fahy et al., 2009). Downstream, we added an extra level between classes and species to enable mapping of lipid identifiers that lack detailed structural information. This concept is also used in the Swiss Lipids system (Aimo et al., 2015). The branch ‘function’ comprises three subcategories: ‘lipid component’ (associated with lipids that are primary regarded as structural component of lipid bilayers), ‘lipid-mediated signaling’ (lipids that have been implicated in signaling) and ‘lipid-storage’ (lipids that are associated with storage, primarily in lipid droplets). In the category ‘cellular localization’, lipid classes that are enriched in particular cellular organelles are linked to their corresponding organelle terms (Holthuis and Menon, 2014; van Meer et al., 2008). The branch ‘physical or chemical properties’ comprises a number of subcategories.

First, a number of chemical descriptions (‘contains fatty acid’, ‘fatty acid unsaturation’, ‘fatty acid length’ and ‘type by bond’) was inferred from the species names. Second, data about ‘intrinsic curvature’ (de Kroon et al., 2013; Thiam et al., 2013) were categorized into either negative, neutral or positive curvature. As data on species-level are limited, curvature was assumed to be predominantly headgroup-dependent and fatty acid composition was neglected. The third subcategory, ‘charge headgroup’, was divided into three groups based on structural data: ‘negative’, ‘positive/zwitter-ion’ and ‘neutral’ (Fahy et al., 2009). This last term comprises also lipids lacking a headgroup. The fourth subcategory in ‘physical or chemical properties’ is 'chain-melting transition temperature'. This property is derived from a number of sources, comprehensively reviewed by Marsh (Marsh, 2010). This dataset covers a range of lipid classes in both glycerophospholipids (PC, PE, PG, PA, PS) and sphingolipids (SM). We made use of multiple regression analysis with lipid class, fatty acid length and unsaturation as predictors to facilitate data extrapolation to previously unreported lipid species. The obtained model was validated by leave-one-out cross-validation (LOOCV). Briefly, one datapoint from the dataset was taken out, after which the model was rebuilt with the remaining points as training set. Subsequently, the selected datapoint was used as validation sample. This procedure was repeated for all the datapoints (**Fig. S2 C**). Next, values predicted by the obtained model of all applicable lipid species present in LION were divided into quintiles with limits based on four reported lipidomics datasets (Andreyev et al., 2010; Haraszti et al., 2016; Köberlin et al., 2015; Lin et al., 2017) and categorized into five representative classes: ‘very low’, ‘low’, ‘average’, ‘high’ or ‘very high’ chain-melting transition temperature (a flow-chart of this procedure is depicted in **Fig. S2 E**).

In addition to these experimental data sets, we also used data (Wassenaar et al., 2015) that was obtained by coarse grain molecular dynamics simulation (MARTINI force-field (Marrink et al., 2004)) and which includes membrane properties ‘bilayer thickness’ and ‘lateral diffusion’. The dataset contains lipids from five common classes of glycerophospholipids (PC, PS, PG, PA, PE), but lacks sphingolipids and sterols. By definition, coarse-grained lipids represent a range of structures. To be able to use the dataset in the ontology system, names of coarse-grained lipids were translated into their representing counterparts. Subsequently, lipid properties were extrapolated to the entire database by multiple regression analysis models (with lipid class, fatty acid length and unsaturation as predictors) and validated by LOOCV (**Fig. S2 A-B**). We followed the same procedure as used for transition temperatures; extrapolated results for both properties were divided into quintiles (based on values of reported datasets (Andreyev et al., 2010; Haraszti et al., 2016; Köberlin et al., 2015; Lin et al., 2017), predicted by our models) and categorized into five representative classes: ‘very low’, ‘low’, ‘average’, ‘high’ or ‘very high’.

The initial structure of LION was build with OBOEdit v.2.3.1 (Wächter and Schroeder, 2010) and formatted as OBO-file. Subsequently, custom R-scripts connected specific terms with more general terms based on the described datasets. The entire ontology can be found as **File S1.**In addition, a table containing all LION-terms with corresponding LION-identifier is provided in **Data S5**.

### Implementation of enrichment analysis tool

To use LION with existing ontology enrichment tools, we used an adapted and generalized version of Bioconductor R-package ‘topGO’ (Alexa and Rahnenfuhrer, 2017). This version, called ‘topOnto’, allows users to include ontologies other than those provided with the package. A custom Perl-script was used to convert the ontology file from OBO to SQLite-format. Apart from this extra feature, the ‘topOnto’ package provides the same functionality as the original version. To perform the enrichment analysis with ‘topOnto’, two statistical approaches were used. In the ‘by target-list mode’, Fisher-exact statistics are used to indicate enrichment. In the ‘by ranking’ mode, Kolmogorov-Smirnov tests are used as ‘global’ statistics. In both approaches, topGO’s classic algorithm was selected. After LION enrichment analysis, raw *P* values were corrected for multiple testing (Benjamini-Hochberg). The R-scripts were used to build the user-friendly web-based tool LION/web (**Note S1**)with R-package ‘shiny’. The application has been made available on the shinyapps.io server as a free online tool, accessible through http://www.lipidontology.com/.

### Cell culture and preparation of fatty acid-albumin complexes

CHO-k1 cells were cultured in Ham’s F-12 medium (Thermo Fisher Scientific, Waltham, MA, USU) supplemented with 7.5% FBS (Thermo Fisher Scientific, Waltham, MA, USU), 100 units/ml penicillin and 100 μg/ml streptomycin (Thermo Fisher Scientific, Waltham, MA, USU). Cells were grown in a humidified incubator at 37°C containing 5% CO_2_ and passaged twice a week. Stocks of 10 mM arachidonic acid, linoleic acid, oleic acid, or palmitic acid (all obtained from Sigma, St. Louis, MO, USA) were complexed to 2 mM fatty-acid free BSA (Sigma, St. Louis, MO, USA), filter-sterilized and stored at −20 °C. All experimental incubations were performed in plastic 6-well culture dishes (Corning, Tewksbury, MA, USA).

### Measuring membrane fluidity

After overnight incubation in the absence or presence of fatty-acids (using fatty acid-free BSA or fatty acids coupled to BSA, respectively), cells were washed and scraped in PBS. Cells were subsequently homogenized on ice with 26-gauge needles (BD Bioscience, San Jose, CA, USA). Homogenates or blanks were mixed with pyrenedecanoic acid (PDA) in the manufacturer’s supplied dilution buffer (Membrane fluidity kit, Abcam, Cambridge, UK) and transferred into a 96-well plate (black plastic with glass bottom, Greiner Bio-One, Frickenhausen, Germany). After 30 minutes of incubation at 37°C, fluorescence spectra (excitation at 360nm, emission between 375-500 nm, 37°C) were measured with a temperature-controlled fluorescence microplate reader (CLARIOstar, BMG Labtech, Offenburg, Germany). Data were processed in R by expressing monomer (370-390 nm) and excimer (470-490 nm) mean fluorescence after blank-subtraction as ratios. Data were expressed as means. Differences were analyzed by two-tailed Student’s t-tests. *P* values < 0.05 were considered significant.

### Lipidomics by LC-MS/MS

After incubation, lipids were extracted as described before (Bligh and Dyer, 1959). Subsequently, lipid extracts were dried under nitrogen and dissolved in 100 μL chloroform/methanol (1:1) and injected (10 μL) on a hydrophilic interaction liquid chromatography (HILIC) column (2.6 μm HILIC 100 Å, 50 × 4.6 mm, Phenomenex, Torrance, CA), eluted by an eluens gradient (flow rate of 1 mL/min) from ACN/acetone (9:1, v/v) to ACN/H_2_O (7:3, v/v) with 10mM ammonium formate, both containing 0.1% formic acid. The column effluent was connected to a heated electrospray ionization (hESI) source of a mass spectrometer (Fusion, Thermo Scientific, Waltham, MA). The measurements were performed with an orbitrap resolution of 120.000, generating 30 data-dependent MS/MS spectra per second in the linear ion trap.

### Lipidomics data analysis

Acquired raw datafiles were converted to mzXML-files and processed with R-package ‘xcms’ v2.99.3 (Smith et al., 2006). After deisotoping, annotation of lipids was performed by matching measured MS-1 m/z values with theoretical m/z values. Lipids with the same or similar m/z values - *e.g.*, BMP(38:4) and PG(38:4) - could by distinguished by differences in retention time (**Fig. S4 and S5**). Lipid annotation containing individual fatty acids as used in **Fig. 2 A** and **Fig. S4** was accomplished by examining MS-2 spectra. When MS-2 spectra were available for a given MS-1 peak, the most abundant fatty acid combination was used to annotate the lipid. The resulting experimental datasets, as well as the public RAW 264.7 macrophage dataset (Andreyev et al., 2010), were normalized by expressing all lipids as ratios of the sum of all intensities per sample. MetaboAnalyst 3.0 (Xia et al., 2015) was used to replace missing values (of the RAW 264.7 dataset) by half of the minimum positive value in the original data, and to perform Principal Component Analysis (with Pareto scaling).

### Software and R-packages

All R-scripts were run with RStudio v1.0.153 (R v3.4.4) with the following packages: ‘shiny’, ‘visNetwork’, ‘data.table’, ‘GMD’, ‘igraph’, ‘reshape2’, ‘ggplot2’, ‘ggthemes’, ‘shinyTree’, ‘shinyWidgets’, ‘shinythemes’, ‘RSQLite’, ‘topOnto’ and ‘xcms’ (Smith et al., 2006). Perl-scripts were run with Perl v5.26.0. All figures were build in R and processed in Cytoscape v3.5.1 or Inkscape v0.92.2.

### Data and code availability

The raw lipidomics data are available as Supplemental Data. The public RAW 264.7 macrophages dataset (Andreyev et al., 2010) is available on the journal’s website. R-package ‘topOnto’ is available at https://github.com/hxin/topOnto, the associated R-package containing the LION database at https://github.com/martijnmolenaar/topOnto.LION2.db.

## REFERENCES

Aimo, L., Liechti, R., Hyka-Nouspikel, N., Niknejad, A., Gleizes, A., Götz, L., Kuznetsov, D., David, F.P.A., van der Goot, F.G., Riezman, H., et al. (2015). The SwissLipids knowledgebase for lipid biology. Bioinforma. Oxf. Engl. 31, 2860–6.

Alexa, A., and Rahnenfuhrer, J. (2017). Gene set enrichment analysis with topGO. Bioconductor.

Andreyev, A.Y., Fahy, E., Guan, Z., Kelly, S., Li, X., McDonald, J.G., Milne, S., Myers, D., Park, H., Ryan, A., et al. (2010). Subcellular organelle lipidomics in TLR-4-activated macrophages. J. Lipid Res. 51, 2785–97.

Ashburner, M., Ball, C.A., Blake, J.A., Botstein, D., Butler, H., Cherry, J.M., Davis, A.P., Dolinski, K., Dwight, S.S., Eppig, J.T., et al. (2000). Gene ontology: tool for the unification of biology. The Gene Ontology Consortium. Nat. Genet. 25, 25–9.

Barry, W.T., Nobel, A.B., and Wright, F.A. (2005). Significance analysis of functional categories in gene expression studies: a structured permutation approach. Bioinforma. Oxf. Engl. 21, 1943–9.

Ben M’barek, K., Ajjaji, D., Chorlay, A., Vanni, S., Forêt, L., and Thiam, A.R. (2017). ER Membrane Phospholipids and Surface Tension Control Cellular Lipid Droplet Formation. Dev. Cell 41, 591–604.e7.

Bigay, J., Gounon, P., Robineau, S., and Antonny, B. (2003). Lipid packing sensed by ArfGAP1 couples COPI coat disassembly to membrane bilayer curvature. Nature 426, 563–6.

Bligh, E.G., and Dyer, W.J. (1959). A rapid method of total lipid extraction and purification. Can. J. Biochem. Physiol. 37, 911–917.

Calder, P.C. (2015). Marine omega-3 fatty acids and inflammatory processes: Effects, mechanisms and clinical relevance. Biochim. Biophys. Acta BBA - Mol. Cell Biol. Lipids 1851, 469–484.

Degtyarenko, K., de Matos, P., Ennis, M., Hastings, J., Zbinden, M., McNaught, A., Alcántara, R., Darsow, M., Guedj, M., and Ashburner, M. (2008). ChEBI: a database and ontology for chemical entities of biological interest. Nucleic Acids Res. 36, D344–50.

Eisinger, J., and Scarlata, S.F. (1987). The lateral fluidity of erythrocyte membranes temperature and pressure dependence. Biophys. Chem. 28, 273–281.

Enkavi, G., Mikkolainen, H., Güngör, B., Ikonen, E., and Vattulainen, I. (2017). Concerted regulation of npc2 binding to endosomal/lysosomal membranes by bis(monoacylglycero)phosphate and sphingomyelin. PLOS Comput. Biol. 13, e1005831.

Fahy, E., Subramaniam, S., Murphy, R.C., Nishijima, M., Raetz, C.R.H., Shimizu, T., Spener, F., van Meer, G., Wakelam, M.J.O., and Dennis, E.A. (2009). Update of the LIPID MAPS comprehensive classification system for lipids. J. Lipid Res. 50 Suppl, S9–14.

Haraszti, R.A., Didiot, M.-C., Sapp, E., Leszyk, J., Shaffer, S.A., Rockwell, H.E., Gao, F., Narain, N.R., DiFiglia, M., Kiebish, M.A., et al. (2016). High-resolution proteomic and lipidomic analysis of exosomes and microvesicles from different cell sources. J. Extracell. Vesicles 5, 32570.

Holthuis, J.C.M., and Menon, A.K. (2014). Lipid landscapes and pipelines in membrane homeostasis. Nature 510, 48–57.

Inda, M.E., Vandenbranden, M., Fernández, A., de Mendoza, D., Ruysschaert, J.-M., and Cybulski, L.E. (2014). A lipid-mediated conformational switch modulates the thermosensing activity of DesK. Proc. Natl. Acad. Sci. U. S. A. 111, 3579–84.

Köberlin, M.S., Snijder, B., Heinz, L.X., Baumann, C.L., Fauster, A., Vladimer, G.I., Gavin, A.C., and Superti-Furga, G. (2015). A Conserved Circular Network of Coregulated Lipids Modulates Innate Immune Responses. Cell 162, 170–183.

de Kroon, A.I.P.M., Rijken, P.J., and De Smet, C.H. (2013). Checks and balances in membrane phospholipid class and acyl chain homeostasis, the yeast perspective. Prog. Lipid Res. 52, 374–94.

Liebisch, G., Vizcaíno, J.A., Köfeler, H., Trötzmüller, M., Griffiths, W.J., Schmitz, G., Spener, F., and Wakelam, M.J.O. (2013). Shorthand notation for lipid structures derived from mass spectrometry. J. Lipid Res. 54, 1523–30.

Lin, L., Ding, Y., Wang, Y., Wang, Z., Yin, X., Yan, G., Zhang, L., Yang, P., and Shen, H. (2017). Functional lipidomics: Palmitic acid impairs hepatocellular carcinoma development by modulating membrane fluidity and glucose metabolism. Hepatol. Baltim. Md 66, 432–448.

Marrink, S.J., de Vries, A.H., and Mark, A.E. (2004). Coarse Grained Model for Semiquantitative Lipid Simulations. J. Phys. Chem. B 108, 750–760.

Marsh, D. (2010). Structural and thermodynamic determinants of chain-melting transition temperatures for phospholipid and glycolipids membranes. Biochim. Biophys. Acta 1798, 40–51.

van Meer, G., Voelker, D.R., and Feigenson, G.W. (2008). Membrane lipids: where they are and how they behave. Nat. Rev. Mol. Cell Biol. 9, 112–24.

Sezgin, E., Levental, I., Mayor, S., and Eggeling, C. (2017). The mystery of membrane organization: composition, regulation and roles of lipid rafts. Nat. Rev. Mol. Cell Biol. 18, 361–374.

Sharpe, H.J., Stevens, T.J., and Munro, S. (2010). A comprehensive comparison of transmembrane domains reveals organelle-specific properties. Cell 142, 158–69.

Smith, C.A., Want, E.J., O’Maille, G., Abagyan, R., and Siuzdak, G. (2006). XCMS: processing mass spectrometry data for metabolite profiling using nonlinear peak alignment, matching, and identification. Anal. Chem. 78, 779–787.

Thiam, A.R., Farese, R.V., and Walther, T.C. (2013). The biophysics and cell biology of lipid droplets. Nat. Rev. Mol. Cell Biol. 14, 775–86.

Wächter, T., and Schroeder, M. (2010). Semi-automated ontology generation within OBO-Edit. Bioinforma. Oxf. Engl. 26, i88–96.

Wassenaar, T.A., Ingólfsson, H.I., Böckmann, R.A., Tieleman, D.P., and Marrink, S.J. (2015). Computational lipidomics with insane: A versatile tool for generating custom membranes for molecular simulations. J. Chem. Theory Comput. 11, 2144–2155.

Xia, J., Sinelnikov, I.V., Han, B., and Wishart, D.S. (2015). MetaboAnalyst 3.0-making metabolomics more meaningful. Nucleic Acids Res. 43, W251–7.

Yang, X., Sheng, W., Sun, G.Y., and Lee, J.C.M. (2011). Effects of fatty acid unsaturation numbers on membrane fluidity and α-secretase-dependent amyloid precursor protein processing. Neurochem. Int. 58, 321–9.

